# PPTC7 limits mitophagy through proximal and dynamic interactions with BNIP3 and NIX

**DOI:** 10.1101/2024.01.24.576953

**Authors:** Lianjie Wei, Mehmet Oguz Gok, Jordyn D. Svoboda, Merima Forny, Jonathan R. Friedman, Natalie M. Niemi

## Abstract

PPTC7 is a mitochondrial-localized PP2C phosphatase that maintains mitochondrial protein content and metabolic homeostasis. We previously demonstrated that knockout of *Pptc7* elevates mitophagy in a BNIP3– and NIX-dependent manner, but the mechanisms by which PPTC7 influences receptor-mediated mitophagy remain ill-defined. Here, we demonstrate that loss of PPTC7 upregulates BNIP3 and NIX post-transcriptionally and independent of HIF-1α stabilization. On a molecular level, loss of *PPTC7* prolongs the half-life of BNIP3 and NIX while blunting their accumulation in response to proteasomal inhibition, suggesting that PPTC7 promotes the ubiquitin-mediated turnover of BNIP3 and NIX. Consistently, overexpression of PPTC7 limits the accumulation of BNIP3 and NIX protein levels in response to pseudohypoxia, a well-known inducer of mitophagy. This PPTC7-mediated suppression of BNIP3 and NIX protein expression requires an intact PP2C catalytic motif but is surprisingly independent of its mitochondrial targeting, indicating that PPTC7 influences mitophagy outside of the mitochondrial matrix. We find that PPTC7 exists in at least two distinct states in cells: a longer isoform, which likely represents full length protein, and a shorter isoform, which likely represents an imported, matrix-localized phosphatase pool. Importantly, anchoring PPTC7 to the outer mitochondrial membrane is sufficient to blunt BNIP3 and NIX accumulation, and proximity labeling and fluorescence co-localization experiments suggest that PPTC7 associates with BNIP3 and NIX within the native cellular environment. Importantly, these associations are enhanced in cellular conditions that promote BNIP3 and NIX turnover, demonstrating that PPTC7 is dynamically recruited to BNIP3 and NIX to facilitate their degradation. Collectively, these data reveal that a fraction of PPTC7 dynamically localizes to the outer mitochondrial membrane to promote the proteasomal turnover of BNIP3 and NIX.

## Introduction

Mitophagy, or mitochondrial-specific autophagy, is a conserved organellar quality control process that promotes the selective turnover of damaged or superfluous mitochondria across cellular conditions (Pickles *et al*, 2018; Uoselis *et al*, 2023). During mitophagy, mitochondria are selectively tagged for degradation by the activation of either ubiquitin-associated pathways or specific mitophagy receptors. Multiple human diseases result from with mutations within mitophagy-associated genes, such as Parkinson Disease (Valente *et al*, 2004; Kitada *et al*, 1998; Zimprich *et al*, 2011) and Amyotrophic Lateral Sclerosis (Maruyama *et al*, 2010). Most of these disease-associated mutations lessen the activation or efficiency of mitophagy, suggesting that enhancing mitophagy may promote mitochondrial function and alleviate various human pathologies (Mishra & Thakur, 2023; Lee & Kim, 2014; Wang *et al*, 2023). Interestingly, however, recent studies show that unrestrained mitophagy may also trigger pathophysiology, particularly through disrupting the regulation of the mitophagy receptors BNIP3 and NIX (Bonnen *et al*, 2013; Gai *et al*, 2013; Cao *et al*, 2023; Elcocks *et al*, 2023; Nguyen-Dien *et al*, 2023).

BNIP3 and NIX have long been associated with mitochondrial turnover. Well-characterized as transcriptional targets of Hypoxia Inducible Factor 1α (HIF-1α), BNIP3 and NIX are upregulated during hypoxia to decrease mitochondrial content due to limited oxygen availability (Ney, 2015). NIX promotes mitochondrial clearance during erythropoiesis to prevent mature red blood cells from consuming the oxygen they carry before it is delivered to distal tissues (Schweers *et al*, 2007). Similarly, BNIP3 and NIX induce mitochondrial turnover and subsequent metabolic reprogramming in various models of cellular differentiation, including neurons (Ordureau *et al*, 2021), myoblasts (Sin *et al*, 2016), and cardiomyocytes (Esteban-Martínez & Boya, 2018; Zhao *et al*, 2020). These studies demonstrate that BNIP3 and NIX can potently decrease mitochondrial content across cell types, indicating the levels of these mitophagy receptors must be tightly regulated to prevent excessive mitochondrial clearance. Consistently, recent studies have shown that loss of the mitochondrial E3 ubiquitin ligase FBXL4 unleashes BNIP3– and NIX-mediated mitophagy, leading to decreased mitochondrial protein levels, mtDNA depletion, and perinatal lethality in mice (Alsina *et al*, 2020; Cao *et al*, 2023; Elcocks *et al*, 2023; Nguyen-Dien *et al*, 2023). Mutations in human *FBXL4* cause Mitochondrial DNA Depletion Syndrome 13 (MTDPS13) a severe pathology characterized by encephalopathy, stunted growth, and metabolic deficiencies (Bonnen *et al*, 2013; Gai *et al*, 2013; Dai *et al*, 2017; Ballout *et al*, 2019). Importantly, these human mutations disrupt the ability of FBXL4 to promote BNIP3 and NIX turnover (Cao *et al*, 2023; Elcocks *et al*, 2023; Nguyen-Dien *et al*, 2023), suggesting that excessive BNIP3 and NIX accumulation constitutes a substantial organismal liability. Despite this, the mechanisms restraining these mitophagy receptors under basal conditions have not been fully defined.

We previously identified the mitochondrial-resident protein phosphatase PPTC7 as a regulator of BNIP3– and NIX-mediated mitophagy (Niemi *et al*, 2023). Knockout of *Pptc7* in mice led to fully penetrant perinatal lethality concomitant to metabolic defects, including hypoketotic hypoglycemia (Niemi *et al*, 2019). Strikingly, tissues and cells isolated from *Pptc7* knockout animals showed robustly decreased mitochondrial protein levels as well as consistently elevated BNIP3 and NIX protein expression (Niemi *et al*, 2019), indicating that unchecked BNIP3 and NIX expression may drive mitochondrial loss through excessive mitophagy. Indeed, knockout of *Bnip3* and *Nix* within the *Pptc7* knockout background largely rescues mitochondrial protein levels and elevated mitophagy (Niemi *et al*, 2023). Additionally, we found that BNIP3 and NIX are hyperphosphorylated in *Pptc7* knockout systems, and that PPTC7 can directly interact with BNIP3 and NIX to facilitate their dephosphorylation in vitro (Niemi *et al*, 2023). These data demonstrate that PPTC7 acts as a critical negative regulator of BNIP3– and NIX-mediated mitophagy. However, the precise molecular mechanism(s) by which PPTC7 influences BNIP3 and NIX protein levels and mitophagic flux remain unclear, particularly given that these proteins reside in separate mitochondrial compartments (Rhee *et al*, 2013; Hung *et al*, 2017). Here, we use a combination of biochemical and cellular assays to demonstrate that PPTC7 proximally and dynamically interacts with BNIP3 and NIX to promote their turnover and limit basal receptor mediated mitophagy.

## Results

### PPTC7 regulates BNIP3 and NIX post-transcriptionally and independent of HIF-1α

We previously reported that BNIP3 and NIX were significantly upregulated in tissues and cells derived from *Pptc7^-/-^* mice (Niemi *et al*, 2019, 2023). Consistently, *Pptc7* knockout (KO) mouse embryonic fibroblasts (MEFs, Figure 1A) and *PPTC7* KO 293T cells (Figure 1B) showed elevated expression of BNIP3 and NIX relative to wild-type cells. To understand the mechanisms underlying BNIP3 and NIX upregulation upon *PPTC7* loss, we investigated the involvement of Hypoxia Inducible Factor-1α (HIF-1α) in mediating this response. *BNIP3* and *NIX* are well-established transcriptional targets of HIF-1α in conditions of hypoxia, as well as pseudohypoxia through various pharmacological stimuli (e.g., the iron chelators deferoxamine (DFO) and deferiprone (DFP) as well as cobalt chloride) (Allen *et al*, 2013; Wang & Semenza, 1993a, 1993b). We thus hypothesized that the increase in BNIP3 and NIX protein levels in *PPTC7* KO cells may be due to elevated HIF-1α activity. To test this, we immunoblotted for HIF-1α expression in wild-type and *PPTC7* KO cells in both basal conditions as well as in the presence of bafilomycin A1 (Baf-A1), a compound previously shown to stabilize HIF-1α (Hubbi *et al*, 2013). These experiments revealed no differences in HIF-1α protein expression between wild-type and *PPTC7* KO cells across tested conditions (Figure 1C). Additionally, we found no significant changes in the abundance of select HIF-regulated proteins relative to other proteins across our previously collected proteomics datasets in *Pptc7* KO mouse tissues (Niemi *et al*, 2019) (Supplemental Figure 1A). We next tested whether BNIP3 and NIX were transcriptionally upregulated but found no significant differences in *BNIP3* or *NIX* mRNA levels between wild-type and *PPTC7* KO 293T cells (Figure 1D). Consistently, we found that the transient transfection of plasmids encoding myc-BNIP3 or FLAG-NIX led to increased protein expression in *PPTC7* KO 293T cells relative to wild-type 293T cells (Supplemental Figures 1B, C). As these plasmids are driven by the same CMV promoter in both wild-type and *PPTC7* KO cells, the elevated BNIP3 and NIX protein expression seen in *PPTC7* KO likely occurs independent of transcriptional changes. Together, these data indicate that PPTC7 influences BNIP3 and NIX protein expression post-transcriptionally and independent of HIF-1α.

**Figure 1:**
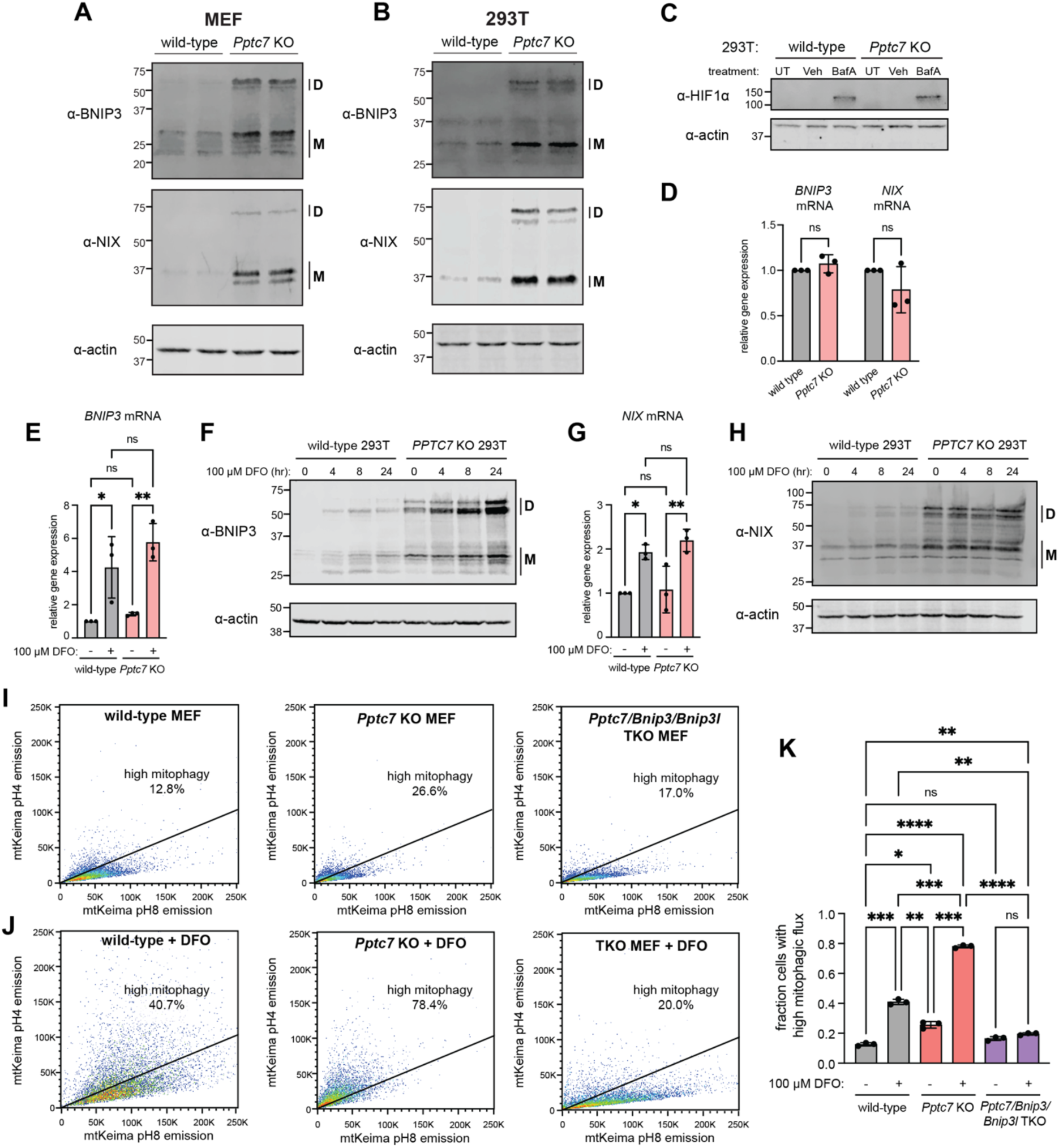
BNIP3 and NIX are upregulated post-transcriptionally and independent of HIF-1α in PPTC7 KO cells. **A**., **B**. Western blots of BNIP3 (top panels), NIX (middle panels) and actin (serving as a load control, bottom panels) in wild-type or *Pptc7* knockout (KO) mouse embryonic fibroblasts (MEF, A.) or in wild-type or *PPTC7* KO 293T cells (B.). “D” indicates dimer species; “M” indicates monomer species. **C**. Western blot for HIF-1α in untreated (UT), vehicle only (veh, 0.2% DMSO) or 100 nM bafilomycin A (BafA) for 16 hours in wild-type and *PPTC7* KO 293T cells. Actin shown as a loading control. **D**. qRT-PCR of BNIP3 and NIX endogenous mRNA levels in wild-type (gray) and *PPTC7* KO (pink) 293T cells. **E**. qRT-PCR of *BNIP3* RNA levels in untreated and DFO-treated (100 μM, 24 hours) wild-type (gray) and *PPTC7* KO (pink) 293T cells. **p<0.01, *p<0.05, ns = not significant, Ordinary one-way ANOVA. Error bars represent standard deviation; data points represent independent experiments. **F**. Western blotting for endogenous BNIP3 levels in wild-type or *PPTC7* KO 293T cells treated with 100 μM DFO for indicated times. Actin shown as a loading control. **G**. qRT-PCR of *NIX* RNA levels in untreated and DFO-treated (100 μM, 24 hours) wild-type (gray) and *PPTC7* KO (pink) 293T cells. **p<0.01, *p<0.05, ns = not significant, Ordinary one-way ANOVA. Error bars represent standard deviation; data points represent independent experiments. **H**. Western blotting for endogenous NIX levels in wild-type or *PPTC7* KO 293T cells treated with 100 μM DFO for indicated times. Actin shown as a loading control. **I**. FACS histograms of basal mitophagy in wild-type (left), *Pptc7* KO (middle) and *Pptc7/Bnip3/Nix* TKO MEFs using the mt-Keima fluorescence assay. Cells undergoing high mitophagy are above the diagonal line; percentages indicated in figure. **J**. FACS histograms of mitophagy rates upon 24 hours of 100 μM DFO treatment in wild-type (left), *Pptc7* KO (middle) and *Pptc7/Bnip3/Nix* TKO MEFs using the mt-Keima fluorescence assay. **K**. Quantification of mt-Keima data shown in I., J. ****p<0.0001, ***p<0.001, **p<0.01, *p<0.05, ns = not significant, Ordinary one-way ANOVA. Error bars represent standard deviation; data points represent individual biological replicates.

The HIF-1α-independent upregulation of BNIP3 and NIX in *PPTC7* KO cells suggests that HIF-1α and PPTC7 regulate BNIP3 and NIX through parallel pathways. If true, we hypothesized that loss of *PPTC7* would not alter BNIP3 and NIX transcriptional induction upon HIF activation, but would promote an additive increase in BNIP3 and NIX protein levels. Indeed, treatment of both wild-type and *PPTC7* KO cells with the iron chelator DFO induced similar fold changes in *BNIP3* (Figure 1E) and *NIX* (Figure 1G) mRNA transcripts across genotypes. At the protein level, expression of BNIP3 (Figure 1F) and NIX (Figure 1H) were substantially higher in DFO-treated *PPTC7* KO cells relative to other tested conditions. Notably, basal protein expression of BNIP3 and NIX in untreated *PPTC7* KO cells exceeded BNIP3 and NIX induction in DFO-treated wild-type cells (Figures 1F and 1H), indicating the magnitude of BNIP3 and NIX upregulation in the absence of *PPTC7*. These experiments further indicate that neither BNIP3 nor NIX is maximally upregulated in *PPTC7* knockout cells in basal conditions.

The additive increase in BNIP3 and NIX expression in *PPTC7* KO cells upon DFO treatment led us to hypothesize that mitophagic flux would be elevated in DFO-treated *PPTC7* knockout cells relative to either condition independently. To test this, we assayed wild-type and *Pptc7* KO MEFs expressing mt-Keima, a pH-sensitive fluorescent mitophagy sensor (Sun *et al*, 2017) using flow cytometry. We first exposed wild-type or *Pptc7* KO MEFs to varying concentrations of DFO and found that only the highest tested dose of DFO, 100 μM, induced mitophagy in wild-type cells (Supplemental Figures 1D-F). Notably, the percentage of cells undergoing high mitophagic flux in wild-type cells treated with 100 μM DFO remained below the rates of mitophagic induction in untreated *Pptc7* KO cells (Supplemental Figures 1D-F). We repeated these experiments in wild-type, *Pptc7* KO, and *Pptc7/Bnip3/Nix* triple knockout (TKO) MEFs and found that 100 μM DFO induced an approximate three-fold increase in mitophagic flux in both wild-type and *Pptc7* KO cells (Figures 1I-K). However, the absolute levels of mitophagy in DFO-treated *Pptc7* KO cells significantly exceeded those of DFO-treated wild-type cells as well as untreated *Pptc7* KO cells (Figures 1I-K). Importantly, mt-Keima-positive *Pptc7/Bnip3/Nix* TKO cells fail to undergo appreciable mitophagy in the presence of 100 μM DFO, demonstrating their necessity in increasing mitophagic flux in *Pptc7* knockout cells (Figures 1I-K). Collectively, these data demonstrate that BNIP3 and NIX are post-transcriptionally upregulated to induce mitophagy in *PPTC7* KO cells, and that transcriptional activation of HIF-1α can substantially enhance BNIP3/NIX protein expression and mitophagy in the absence of PPTC7.

### PPTC7 enables BNIP3 and NIX turnover through proteasomal degradation

The elevated protein expression of BNIP3 and NIX in *PPTC7* knockout cells implies that PPTC7 alters the synthesis or turnover rates of these mitophagy receptors. BNIP3 and NIX turnover has emerged as a critical regulatory step in limiting basal mitophagy, as evidenced by recent studies on the E3 ubiquitin ligase FBXL4 (Cao *et al*, 2023; Elcocks *et al*, 2023; Nguyen-Dien *et al*, 2023). Loss of *FBXL4* phenotypically mirrors *PPTC7* KO in mice, and *PPTC7* and *FBXL4* have significant and positively correlated essentiality profiles across over one-thousand cancer cell lines (Supplemental Figure 2A, (Dempster *et al*, 2019)). These data suggest that PPTC7 and FBXL4 influence BNIP3 and NIX via similar mechanisms, leading us to hypothesize that BNIP3 and NIX have decreased turnover rates in *PPTC7* knockout cells relative to wild-type cells.

To test this, we first sought to identify an experimental condition in which we could quantify endogenous BNIP3 and NIX turnover. We noted that DFO-mediated iron chelation was previously shown to decrease the protein level of select mitochondrial proteins in a manner that was reversible upon compound washout (Rensvold *et al*, 2013). As BNIP3 and NIX accumulate in response to DFO (Figure 1), we hypothesized that washout of DFO would induce BNIP3 turnover due to its short half-life (Schäfer *et al*, 2022) (Figure 2A). Indeed, treatment of wild-type 293T cells with DFO increased BNIP3 levels in a time-dependent manner, which returned to at or near baseline (i.e., untreated) levels 24 hours after DFO washout (Figure 2B). Thus, DFO washout constitutes an experimental system in which we could test the effects of PPTC7 on the turnover of endogenous BNIP3. We repeated these experiments in wild-type and *PPTC7* knockout 293T cells and found that BNIP3 and NIX exhibited blunted turnover in *PPTC7* KO cells (Figures 2C-F, Supplemental Figure 2B-E). Modeling of BNIP3 and NIX decay rates showed that loss of PPTC7 extends the half-life of monomeric and dimeric populations of BNIP3 and NIX (Figures 2C-F). While the dimer populations of BNIP3 and NIX have at least a doubling of protein half-life in *PPTC7* KO cells relative to wild-type cells, the half-lives of the monomeric populations of each mitophagy receptor could not be effectively modeled in *PPTC7* KO cells, consistent with substantial suppression of the turnover of these species of BNIP3 and NIX in the absence of PPTC7 (Figures 2C-F).

**Figure 2:**
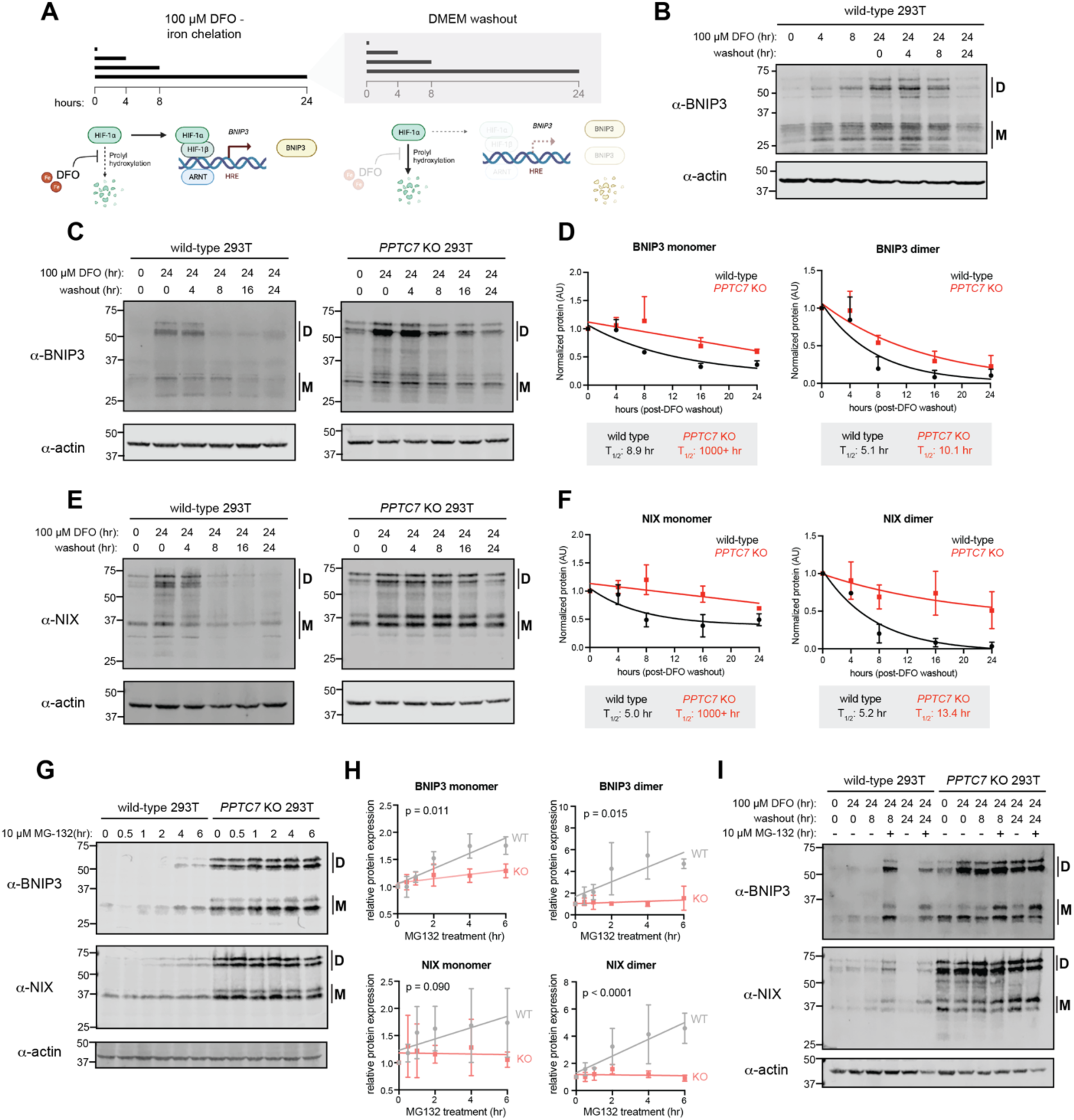
BNIP3 and NIX have decreased turnover rates and are less responsive to proteasomal inhibition in PPTC7 KO cells. **A.** Schematic of DFO treatment and washout timeline and mechanism. **B**. Western blot of endogenous BNIP3 protein after indicated times of DFO treatment and washout, when applicable. “D” indicates dimer species; “M” indicates monomer species. Actin shown as a loading control. **C**. Western blot of endogenous BNIP3 protein after indicated times of DFO treatment and washout in wild-type (left panel) and *PPTC7* KO (right panel) 293T cells. “D” indicates dimer species; “M” indicates monomer species. Actin shown as loading control. **D**. Quantification of data shown in C. BNIP3 monomer (left graph) or dimer (right graph) bands were quantified using densitometry, averaged, and plotted over time. Data were fit with a one phase decay model to calculate protein half-lives (T_1/2_), which are shown below each graph. Error bars represent standard deviations of normalized densitometry across three independent experiments. **E**. Western blot of endogenous NIX protein after indicated times of DFO treatment and washout in wild-type (left panel) and *PPTC7* KO (right panel) 293T cells. “D” indicates dimer species; “M” indicates monomer species. Actin shown as loading control. **F**. Quantification of data shown in E. NIX monomer (left graph) or dimer (right graph) bands were quantified using densitometry, averaged, and plotted over time. Data were fit with a one phase decay model to calculate protein half-lives (T_1/2_), which are shown below each graph. Error bars represent standard deviations of normalized densitometry across three independent experiments. **G**. Western blots of endogenous BNIP3 (top panel) and NIX (bottom panel) in wild-type and *PPTC7* KO 293T cells upon treatment with 10 μM MG-132 for the indicated timeframes. “D” indicates dimer species; “M” indicates monomer species. Actin shown as a loading control. **H**. Quantification of BNIP3 and NIX monomer (left graphs) and dimer (right graphs) populations in wild-type (gray) and *PPTC7* KO (pink) cells shown in G. Bands were quantified using densitometry, averaged, and plotted over time. Data were analyzed via linear regression, and significance between slopes was calculated using Analysis of Covariance (ANCOVA). **I**. Western blot of endogenous BNIP3 and NIX proteins in wild-type and *PPTC7* KO cells after DFO treatment and subsequent washout in the presence or absence of 10 μM MG-132. “D” indicates dimer species; “M” indicates monomer species. Actin shown as a loading control.

Given the slowed rates of BNIP3 and NIX turnover in *PPTC7* KO cells, we sought to understand the pathway(s) contributing to their degradation. The phenotypic similarities between loss-of-function models of *PPTC7* and *FBXL4*, as described above, suggest that these two proteins function in a similar pathway. Additionally, previous work has shown that BNIP3 accumulates in response to the proteasomal inhibitor MG-132 (Park *et al*, 2013; Poole *et al*, 2021). If knockout of *PPTC7* suppresses the proteasomal degradation of BNIP3 and NIX, we hypothesized that *PPTC7* KO cells would be less responsive to the proteasomal inhibitor MG-132 than matched wild-type cells. We found that BNIP3 and NIX levels increased in wild-type cells upon MG-132 treatment (Figures 2G, H), but that the level of each receptor was significantly less responsive to MG-132 in *PPTC7* KO cells (Figures 2G, H). To further test this model, we exploited the DFO washout assay, predicting that if BNIP3 and NIX were turned over by proteasomal degradation upon DFO washout, MG-132 treatment would slow their turnover rates in wild-type cells but have a diminished effect in *PPTC7* KO cells. We found that, upon DFO washout, MG132 treatment causes BNIP3 and NIX accumulation in wild-type cells, while the levels of these proteins remain largely unchanged in *PPTC7* KO cells under identical conditions (Figure 2I). Overall, these data demonstrate that PPTC7 enables the efficient turnover of BNIP3 and NIX in a manner that largely depends upon proteasomal degradation.

### PPTC7 requires an intact active site but not a mitochondrial targeting sequence to limit BNIP3 and NIX accumulation

As loss of *PPTC7* increases BNIP3 expression, we hypothesized that PPTC7 overexpression may diminish BNIP3 upregulation induced by pseudohypoxia. To test this, we overexpressed either cytosolic GFP or PPTC7-GFP in HeLa cells that were treated with the pseudohypoxia inducer cobalt chloride (CoCl_2_). We fixed and immunolabeled these cells to examine endogenous BNIP3 levels relative to the general mitochondrial marker TOMM20. CoCl_2_ robustly upregulated BNIP3 protein expression, and mitochondrial BNIP3 staining could be detected in nearly all cells transfected with cytosolic GFP (Figure 3A). In contrast to cytosolic GFP, PPTC7-GFP co-localized with the mitochondrial marker TOMM20 as expected (Figure 3A). Notably, mitochondrial BNIP3 signal was rarely detected in the subset of cells expressing PPTC7-GFP (Figures 3A, B). On a functional level, overexpression of PPTC7 additionally suppressed CoCl_2_-induced mitophagic flux relative to identically treated wild-type mt-Keima cells (Figure 3C, Supplemental Figure 3A). Finally, overexpression of PPTC7 in *Pptc7* KO MEFs rescued basal BNIP3 protein expression to levels seen in wild-type MEFs (Supplemental Figure 3B). Collectively, these data show that overexpressed PPTC7 limits BNIP3 protein expression and mitophagy induction in both wild-type and *Pptc7* KO cells.

**Figure 3:**
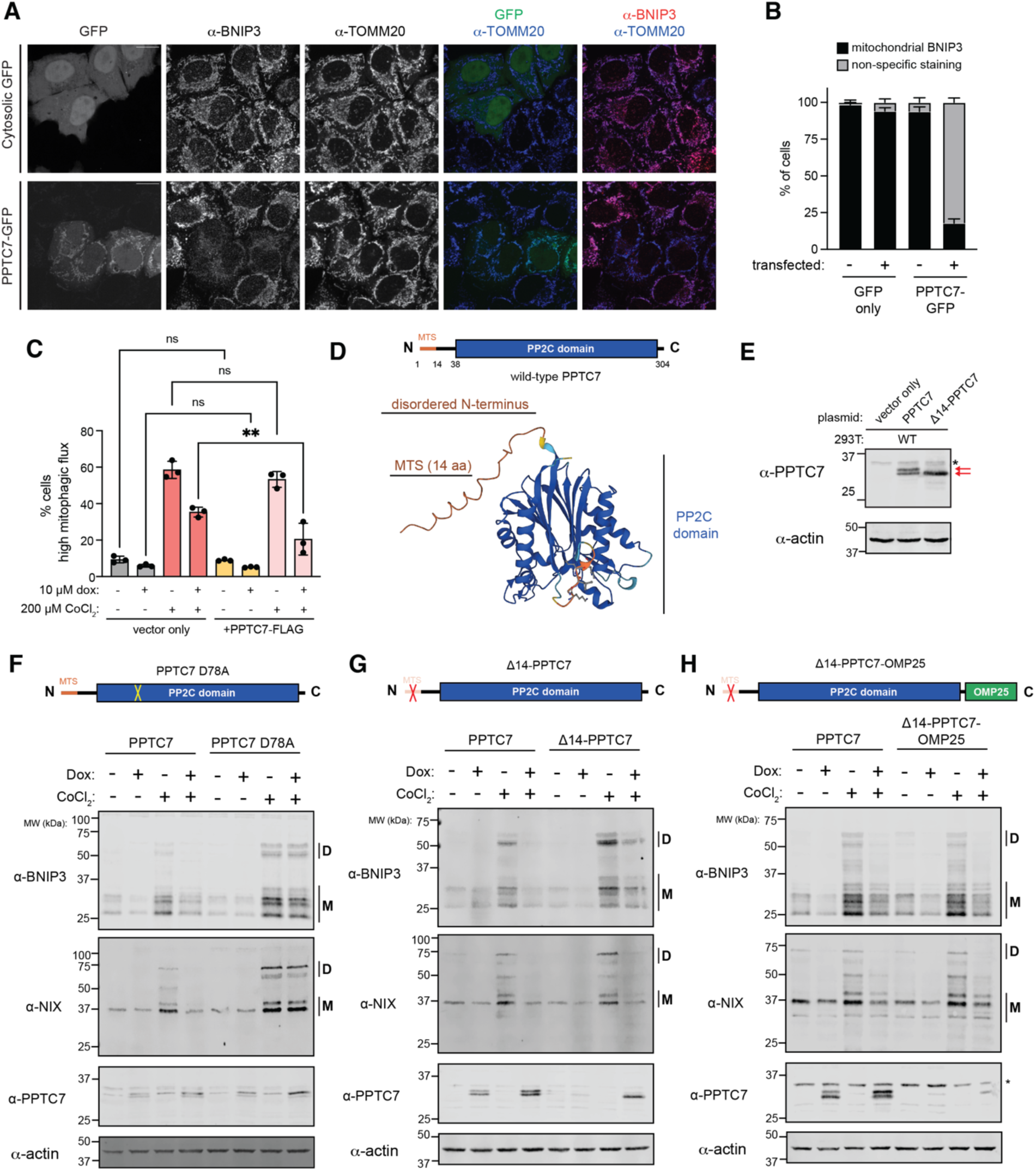
PPTC7 requires an intact catalytic motif but not mitochondrial targeting to suppress BNIP3 and NIX accumulation. **A**. Representative single plane confocal images of GFP only (top panels) or PPTC7-GFP (bottom panels) expressed in HeLa cells treated for 12 hours with 500 μM CoCl_2_. Cells were stained for BNIP3 (second column) or TOMM20 (third column). Overlays are shown for GFP and TOMM20 (fourth column) and GFP and BNIP3 (fifth column). Scale bar = 20 μm. **B**. Quantification of data shown in A, mitochondrial BNIP3 staining versus non-specific staining in cells overexpressing GFP only or PPTC7-GFP versus matched untransfected controls for each experiment. Error bars represent standard error of the mean of three independent experiments. **C**. Quantification of mt-Keima positive mitophagic flux in HeLa FLP-IN TRex cells expressing vector only (left) or PPTC7-FLAG in the presence of 10 μM doxycycline (dox, to promote PPTC7 expression), 200 μM cobalt chloride (CoCl_2_), or both. **p<0.01, ns = not significant, ordinary one-way ANOVA. Error bars represent standard deviation. Each dot represents an independent biological replicate. **D**. Schematic of PPTC7 features, including a mitochondrial targeting sequence (MTS) and PP2C phosphatase domain, top. Bottom, AlphaFold2 representation of PPTC7 structure; predicted disordered N-terminus and PP2C phosphatase domains indicated. **E**. Western blot of 293T cells overexpressing PPTC7 or a ΔMTS-PPTC7 mutant. Red arrows indicate dual species in wild-type PPTC7 expression. * represents a non-specific band. Actin shown as a loading control. **F**.-**H**. Western blots of BNIP3, NIX, and PPTC7 depicting the ability of various mutants of PPTC7 to suppress BNIP3 and NIX accumulation in response to CoCl2 treatment. “D” indicates dimer species, “M” indicates monomer species. In F, a catalytic mutant of PPTC7, D78A, cannot effectively suppress BNIP3 and NIX accumulation relative to wild-type PPTC7. In G, the ΔMTS-PPTC7 mutant partially or fully suppresses BNIP3 and NIX accumulation, respectively. In H, a mutant that artificially anchors PPTC7 to the outer mitochondrial membrane, ΔMTS-PPTC7-OMP25, suppresses BNIP3 and NIX accumulation. Actin shown as a loading control.

As these experiments indicated that overexpressed PPTC7 was properly localized and functional, we generated a series of mutants to understand the mechanism by which PPTC7 limits BNIP3 and NIX accumulation. PPTC7 consists of a PP2C phosphatase domain that is preceded by a predicted 38-residue disordered N-terminus (Figure 3C), which includes a mitochondrial targeting sequence (i.e., MTS) that is processed at amino acid 14 (Calvo *et al*, 2017). Interestingly, when western blotting for overexpressed PPTC7, we found that the protein ran as a doublet, while a ΔMTS-PPTC7 mutant ran at the same molecular weight as the bottom band (Figure 3D), suggesting that wild-type PPTC7 expressed as both a full-length and processed form. Given these molecular insights, we used this overexpression system to test the necessity of PPTC7 catalytic activity and mitochondrial targeting in suppressing BNIP3 and NIX induction under pseudohypoxia. We first mutated the PP2C phosphatase domain of PPTC7 at a key catalytic residue, D78.

We previously demonstrated that recombinant PPTC7 D78A was unable to dephosphorylate BNIP3 and NIX on mitochondria isolated from *Pptc7* KO MEFs (Niemi *et al*, 2023). Consistently, recombinant wild-type PPTC7, but not D78A PPTC7, caused a collapse in the laddering pattern of monomeric BNIP3 in mitochondria isolated from wild-type and *PPTC7* knockout 293T cells (Supplemental Figure 3C). These data show that not only that PPTC7 D78A lacks catalytic activity, but also that the upper monomeric bands seen on BNIP3 western blots represent phosphorylated intermediates that can be directly dephosphorylated by PPTC7. While the D78A mutant lacks phosphatase activity, mutation of PPTC7 D78 may also disrupt its physical interaction with BNIP3 or NIX, as AlphaFold2 multimer models indicate that BNIP3 and NIX dock to PPTC7 proximal to D78 (Supplemental Figures 3D,E). Via either mechanism, we predicted that a D78A mutant would be unable to suppress BNIP3 and NIX accumulation during pseudohypoxia. We overexpressed wild-type and D78A PPTC7 in HeLa FLP-IN T-REx cells and found that both constructs were doxycycline-induced to similar extents and, interestingly, also expressed at higher levels in the presence of CoCl_2_, similar to BNIP3 and NIX (Figure 3F). While overexpression of wild-type PPTC7 decreased BNIP3 and NIX accumulation in response to CoCl_2_ treatment, the D78A mutant failed to suppress the induction of these mitophagy receptors (Figure 3F), indicating that PPTC7 requires an intact catalytic motif to influence BNIP3 and NIX.

We next examined whether disrupting the mitochondrial localization of PPTC7 would affect its ability to suppress CoCl_2_-induced BNIP3 and NIX levels. We overexpressed the ΔMTS-PPTC7 construct HeLa FLP-IN TREx cells, hypothesizing that it would fail to influence CoCl_2_-induced BNIP3 and NIX expression due to its inability to target to mitochondria. Surprisingly, we found that ΔMTS-PPTC7 fully suppressed CoCl_2_-mediated NIX accumulation, and partially suppressed BNIP3 accumulation (Figure 3G). This rescue of BNIP3 and NIX expression is consistent with a model in which a non-targeted PPTC7 can influence mitophagy at sufficient (i.e., overexpressed) levels. This, in combination with the fact that wild-type PPTC7 runs as a doublet, suggested that PPTC7 may exist in two populations: one outside of mitochondria (e.g., a full length, unprocessed form) and one that localizes to the mitochondrial matrix (e.g., ΔMTS-PPTC7, which lacks its mitochondrial targeting sequence upon processing after import). If true, full length PPTC7 may reside outside of mitochondria prior to its import, placing it proximal to BNIP3 and NIX. We thus hypothesized that artificially anchoring PPTC7 to the outer mitochondrial membrane (OMM) would block BNIP3 and NIX accumulation in response to pseudohypoxia. We engineered a ΔMTS-PPTC7-OMP25 construct that both lacks an MTS and is fused to OMP25, a C-terminal tail anchored protein that targets to the OMM (Horie *et al*, 2002). Consistent with our hypothesis, ΔMTS-PPTC7-OMP25 blunts the accumulation of BNIP3 and NIX under both basal and CoCl_2_-treated conditions (Figure 3H), suggesting an OMM-localized pool of PPTC7 influences BNIP3 and NIX protein expression. Collectively, these data show that PPTC7 requires an intact active site but not its mitochondrial targeting sequence to suppress BNIP3 and NIX expression in pseudohypoxic conditions, consistent with a role for PPTC7 in regulating these mitophagy receptors outside of mitochondria.

### PPTC7 proximally and dynamically interacts with BNIP3 and NIX in cells

Our data demonstrating that anchoring PPTC7 to the OMM blunts BNIP3 and NIX accumulation (Figure 3H) indicates that a pool of PPTC7 may reside outside of mitochondria to directly interact with BNIP3 and NIX to promote their proteasomal turnover. We sought to test this model by determining whether BNIP3 and NIX interact with PPTC7 in cells through miniTurbo-based proximity labeling experiments. We expressed PPTC7-V5-miniTurbo, as well as two control constructs, in 293T cells with or without DFO treatment. We treated half of the samples with exogenous biotin (which facilitates miniTurbo-based proximity labeling) and left the remaining cells untreated to control for potential non-specific interactions. After biotinylation, we lysed cells, performed a pulldown with streptavidin beads to enrich for biotinylated proximal interactors, and probed for an interaction with BNIP3 and NIX via western blot. While V5-PPTC7-miniTurbo expressed equally across all conditions, the streptavidin pulldown revealed interactions with BNIP3 and NIX only in the biotin-treated samples, demonstrating specific proximal labeling (Figure 4A). Importantly, these interactions were also specific to PPTC7-V5-miniTurbo and were not seen in either vector only or V5-miniTurbo control samples (Figure 4A). These data, combined with our data demonstrating that recombinant PPTC7 can directly dephosphorylate BNIP3 in vitro (Supplemental Figure 2D), strongly suggest that PPTC7 directly interacts with BNIP3 and NIX within the native cellular environment.

**Figure 4:**
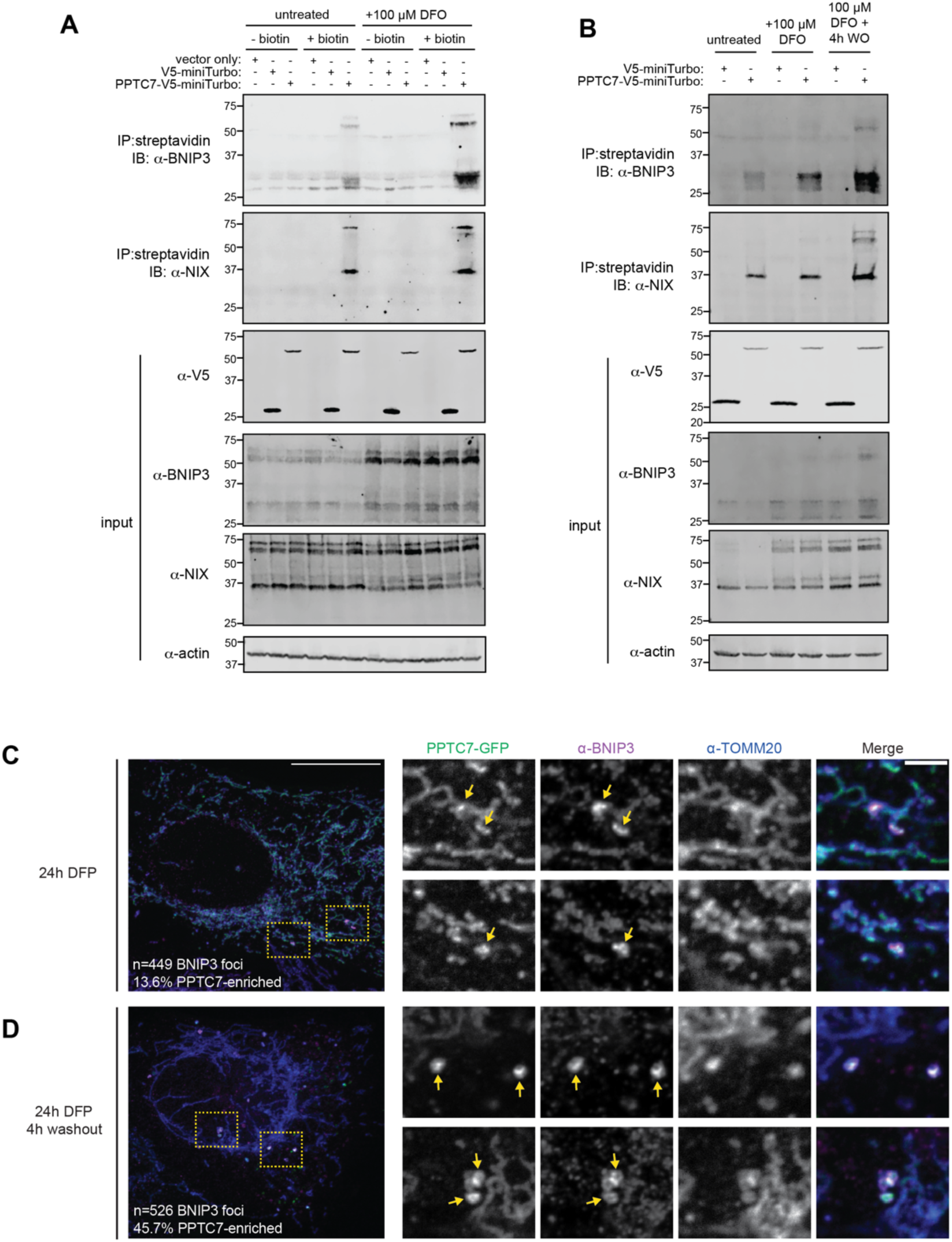
PPTC7 proximally and dynamically interacts with BNIP3 and NIX in cells. **A**. Proximity labeling of PPTC7-V5-miniTurbo in 293T cells with or without 24-hour deferoxamine (DFO) treatment. PPTC7-V5-miniTurbo, as well as vector only or V5-miniTurbo only controls, were transfected into 293T cells. Streptavidin pulldowns were used to enrich for PPTC7-V5-miniTurbo interactors, which were run on SDS-PAGE gels and western blotted for BNIP3 (top blot) or NIX (2^nd^ blot). Only PPTC7-V5-miniTurbo + biotin samples pulled down BNIP3 and NIX (lanes 6 and 12, streptavidin pull down gels), indicating specific binding. Western blots shown for reaction input for pulldowns for V5 (showing miniTurbo constructs), BNIP3, NIX, and actin (serving as a load control). **B**. Proximity labeling of PPTC7-V5-miniTurbo in 293T cells with after 24-hour DFO treatment with or without 4 hour DFO washout. Streptavidin pulldowns were used to enrich for PPTC7-V5-miniTurbo interactors as described in A. Western blots shown for reaction input for pulldowns as described in A. **C**. A representative maximum z-projection confocal image (left) and corresponding single plane insets (right) are shown of a U2OS cell overexpressing PPTC7-GFP and treated with deferiprone (DFP) for 24 hours. Cells were fixed and stained for BNIP3 and TOMM20 to visualize co-enrichment of PPTC7 with BNIP3-enriched foci (n=449). **D**. As in C for a cells treated for 24h with DFP and washed for an additional 4h prior to fixation. Cells were stained to visualize co-enrichment of PPTC7 with BNIP3-enriched foci (n=526).

These data support a model in which PPTC7 is directly recruited to BNIP3 and NIX to promote their turnover. Our DFO washout experiments offer insights into the kinetics of this turnover, which allowed us to test whether the PPTC7-BNIP3/NIX interactions are dynamic throughout the turnover process. Our experiments show that BNIP3 is present at near peak levels 4 hours post-DFO washout (Figures 2B, C), leading us to hypothesize that PPTC7 would have enhanced recruitment to BNIP3 and NIX during acute DFO washout to facilitate the turnover of BNIP3 and NIX. To test this, we repeated proximity labeling of PPTC7-V5-miniTurbo under conditions of acute (i.e., 4 hour) DFO washout. Interestingly, while the PPTC7-BNIP3/NIX interactions are enriched upon 24 hours of DFO treatment, similar to our previous experiment, they are further enhanced upon acute 4-hour DFO washout (Figure 4B). These data show that PPTC7 is dynamically recruited to BNIP3 and NIX to promote their turnover upon resolution of DFO-mediate pseudohypoxia.

To further explore the dynamic nature of the PPTC7-BNIP3 interaction, we exploited the recent observation that BNIP3 enriches at LC3-positive punctate structures which likely represent nascent mitophagosomes (Gok *et al*, 2023). We hypothesized that PPTC7 would also enrich at these punctate structures under conditions of pseudohypoxia, and that this localization would be enriched upon resolution of pseudohypoxia. We overexpressed PPTC7-GFP in U2OS cells treated with the iron chelator deferiprone (DFP) for 24 hours, and then fixed and immunolabeled for BNIP3 and TOMM20. Examination of over 400 BNIP3-enriched foci in DFP-treated cells revealed that ∼14% of these structures were co-enriched for PPTC7-GFP (Figure 4C). Importantly, these foci were co-localized with the mitochondrial marker TOMM20 (Figure 4C), demonstrating these interactions occur at mitochondria and not other organelles, such the ER, where BNIP3 has been reported to localize (Zhang *et al*, 2009; Hanna *et al*, 2012). Remarkably, we found that PPTC7-GFP showed almost a three-fold increase co-enrichment (∼46%) with BNIP3-enriched foci 4 hours after DFP washout relative to DFP treatment alone (Figure 4D). These data, along with our proximity labeling experiments, suggest that PPTC7 is actively recruited to BNIP3 and NIX under conditions that promote their turnover. Collectively, our data are consistent with a model in which a pool of PPTC7 dynamically localizes outside of mitochondria to associate with BNIP3 and NIX–likely through a direct interaction–to promote their ubiquitin-mediated turnover.

## Discussion

PPTC7 is one of twelve phosphatases that localize to mammalian mitochondria (Niemi & Pagliarini, 2021). Conserved through budding yeast (where it is named Ptc7p), PPTC7 has been linked to the maintenance of metabolism and mitochondrial homeostasis across organisms (Martín-Montalvo *et al*, 2013; Guo *et al*, 2017a, 2017b; Gonzalez-Mariscal *et al*, 2017; González-Mariscal *et al*, 2018; Niemi *et al*, 2019, 2023). While we previousy have performed phosphoproteomics to identify candidate substrates of Ptc7p (Guo *et al*, 2017a, 2017b) and PPTC7 (Niemi *et al*, 2019, 2023), the precise roles of these phosphatases in regulating mitochondrial homeostasis remain unclear. In this study, we begin to illuminate important aspects underlying the regulation and function of PPTC7 and how this phosphatase influences BNIP3– and NIX-mediated mitophagy.

The data presented in this study demonstrate that BNIP3 and NIX turnover is a tightly regulated and highly dynamic process. Using a model of DFO-mediated iron chelation followed by compound washout, we show that endogenous BNIP3 and NIX rapidly turn over in wild-type cells in a manner that is slowed by proteasomal inhibition. Our data further show that BNIP3 and NIX have longer half-lives and are less responsive to proteasomal inhibition in *PPTC7* KO cells, consistent with a model in which the phosphatase functions to promote the ubiquitin-mediated turnover of these mitophagy receptors. Indeed, a study published during the preparation of this manuscript showed that PPTC7 coordinates BNIP3 and NIX degradation by acting as a molecular scaffold for the E3 ligase FBXL4 (Sun *et al*, 2023) at the outer mitochondrial membrane. Consistently, we find that PPTC7 does not require its mitochondrial targeting sequence to suppress BNIP3 and NIX accumulation, and anchoring PPTC7 on the outer mitochondrial membrane decreases BNIP3 and NIX protein levels under pseudohypoxia. Furthermore, we find that PPTC7 associates with BNIP3 and NIX via proximity labeling experiments, demonstrating a likely direct interaction between the phosphatase and these mitophagy receptors in cells. These data indicate that PPTC7 has meaningful functions at the outer mitochondrial membrane to regulate BNIP3– and NIX-mediated mitophagy.

The apparent dual functionality of PPTC7 across mitochondrial compartments leads to interesting questions regarding its regulation. The data in this study indicate that PPTC7 is actively recruited to BNIP3 and NIX to promote their degradation, as the proximal interactions between PPTC7 and BNIP3/NIX are enriched upon resolution of pseudohypoxia (i.e., the washout of iron chelator). This conclusion is further supported by the enhanced colocalization of fluorescently tagged PPTC7 with BNIP3-positive punctae in cells in conditions of iron chelator washout. These data suggest not only a model of dynamic recruitment, but also a precisely coordinated spatial organization of BNIP3 and PPTC7 at foci that likely represent mitophagosome formation sites (Gok *et al*, 2023). Interestingly, TMEM11 was also found to co-localize with BNIP3 and NIX at mitophagic punctae, and its knockout increases both BNIP3-positive foci as well as mitophagic flux (Gok *et al*, 2023). These data suggest that PPTC7 and TMEM11 may function in similar complexes to regulate BNIP3 and NIX mediated turnover and/or mitophagy. Whether these complexes include FBXL4 and how they may actively remodel to promote or limit mitophagy are key questions regarding the dynamic regulation of BNIP3– and NIX-mediated mitophagy.

Previous studies of *Fbxl4* and *Pptc7* knockout mouse models show that each display similar pathophysiological profiles, including metabolic dysfunction, broad loss of mitochondrial protein levels, and perinatal lethality. Importantly, knockout of *Bnip3* and *Nix* alleviates many of these phenotypes in cells (Cao *et al*, 2023; Elcocks *et al*, 2023; Nguyen-Dien *et al*, 2023; Niemi *et al*, 2023) and mice (Cao *et al*, 2023; Sun *et al*, 2023), suggesting that loss of *Fbxl4* or *Pptc7* drives excessive BNIP3– and NIX-mediated mitophagy. Despite these advances, the precise mechanism by which PPTC7 influences BNIP3 and NIX protein levels remain unclear. We previously found that BNIP3 and NIX are hyperphosphorylated in *PPTC7* knockout cells and tissues, suggesting PPTC7 may influence mitophagy via their dephosphorylation (Niemi *et al*, 2019, 2023). Indeed, phosphorylation of BNIP3 and NIX can enhance their stability or their ability to induce mitophagy (He *et al*, 2022; Poole *et al*, 2021; Zhu *et al*, 2013; Rogov *et al*, 2017), suggesting dephosphorylation may be required to suppress their activity. However, Sun et al. suggest that PPTC7 promotes BNIP3 and NIX degradation independent of its phosphatase activity, as retains its ability to limit BNIP3 and NIX accumulation in the presence of cadmium, a PP2C phosphatase inhibitor (Sun *et al*, 2023). Interestingly, however, their data demonstrate that an inactive PPTC7 mutant (D78A/G79A) cannot suppress BNIP3 and NIX accumulation or mitophagy induction, which is consistent with our data demonstrating that D78A PPTC7 neither dephosphorylates BNIP3 in vitro, nor blocks BNIP3 and NIX accumulation in conditions of pseudohypoxia (Figure 3). One possible explanation for this discrepancy is that PPTC7 may directly bind to BNIP3 and NIX within its active site, consistent with Alphafold multimer models (Supplemental Figure 3). If so, mutation of the PPTC7 active site could disallow BNIP3 and NIX binding independent of its catalytic activity. Alternatively, it is possible that cadmium does not completely block PPTC7 in cells, allowing low level phosphatase activity that can facilitate BNIP3 and NIX degradation. Careful dissection of the mechanisms by which the D78A mutation PPTC7 limits BNIP3– and NIX-mediated mitophagy, as well as how phosphorylation affects BNIP3 and NIX stability or activity should be active areas of investigation in the future.

Collectively, our data reveal an unexpected role for the mitochondrial protein phosphatase PPTC7 in the regulation of receptor-mediated mitophagy at the outer mitochondria membrane. While knockout of BNIP3 and NIX largely rescue the decreases in mitochondrial protein levels present in *Pptc7* KO tissues and cells, their collective knockout fails to fully rescue metabolic defects seen in *Pptc7* KO cells (Niemi *et al*, 2023). This, together with our data indicating PPTC7 exists in two distinct pools in cells, suggests that PPTC7 regulates additional mitochondrial functions within the matrix. It is possible that PPTC7 mediates such functions by influencing the mitophagic selectivity of matrix-localized substrates, as occurs in yeast (Tal *et al*, 2007; Abeliovich *et al*, 2013; Kolitsida *et al*, 2019, 2023). Alternatively, it is possible that PPTC7 maintains mitochondrial metabolism completely independent of its role in mitophagic signaling, potentially through the regulation of one or more candidate substrates previously identified via phosphoproteomic analyses (Niemi *et al*, 2019, 2023). Disentangling the matrix-mediated functions of PPTC7 versus those that regulate functions outside of mitochondria, and how these ultimately relate to mitophagic flux, will be a key next step in understanding the role of this phosphatase in modulating mitochondrial homeostasis.

## Acknowledgements

We thank Edrees Rashan, Keshav Kailash, Michael McKenna, and the Niemi Laboratory for careful reading of the manuscript and for helpful discussions on this work. We thank Julia Pagan and Keri-Lyn Kozul (University of Queensland, Australia) for key resources and for important discussions related to this manuscript. We thank the Flow Cytometry and Fluorescence-Activated Cell Sorting Core at Washington University School of Medicine for equipment and support for the flow cytometry experiments. This work was supported by R35GM151130 (to N.M.N.) and R35GM137894 (to J.R.F.). The UT Southwestern Quantitative Light Microscopy Facility, which is supported in part by NIH P30CA142543, provided access to the Nikon SoRa microscope (purchased with NIH 1S10OD028630-01). L.W. was supported by the MilliporeSigma Predoctoral Fellowship in Honor of Dr. Gerty T. Cori at Washington University. J.S. was supported through the Summer Undergraduate Research Program put on within the Department of Biochemistry and Molecular Biophysics at Washington University School of Medicine in St. Louis.

---

**Supplemental Figure 1.**
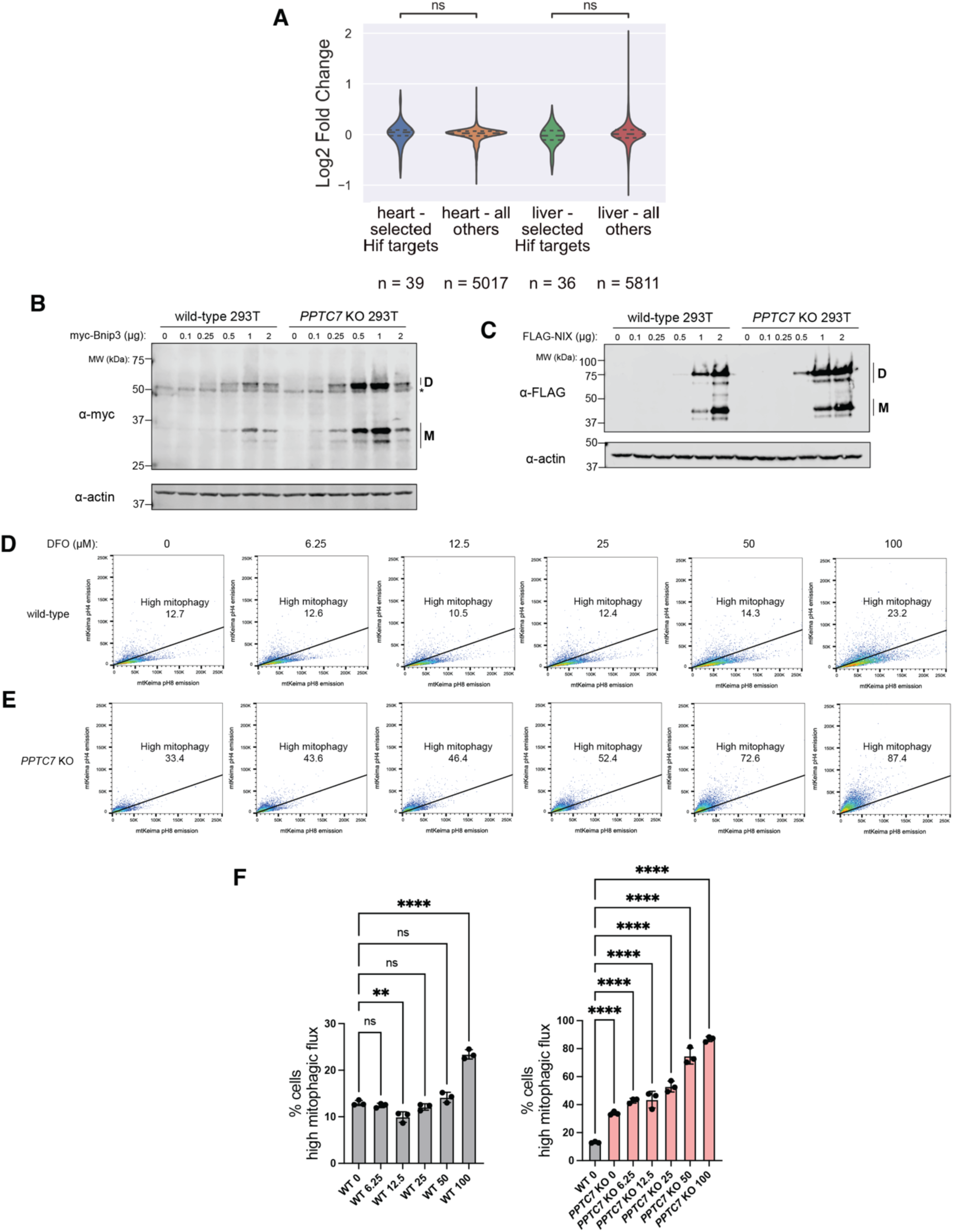
**A**. Analysis of select HIF responsive targets in *Pptc7* KO heart and liver proteomics datasets collected in Niemi et al., Nat Commun, 2019. **B**.,**C**. Western blots of exogenous BNIP3 (B) or NIX (C) expressed in wild-type of *PPTC7* KO 293T cells at various plasmid concentrations. “D” indicates dimer species, “M” indicates monomer species. Actin shown as a loading control. **D**., E. Histograms of FACS analysis of mt-Keima positive wild-type (D) and *Pptc7* KO (E) MEFs showing high mitophagy rates in response to variable DFO concentrations. **F**. Statistical analysis of data shown in D., E., grey bars = wild-type samples, pink bars = *Pptc7* KO samples. ****p<0.0001, ***p<0.001, **p<0.01, ns = not significant, ordinary one-way ANOVA. Error bars represent standard deviation, data points represent individual biological replicates.

**Supplemental Figure 2:**
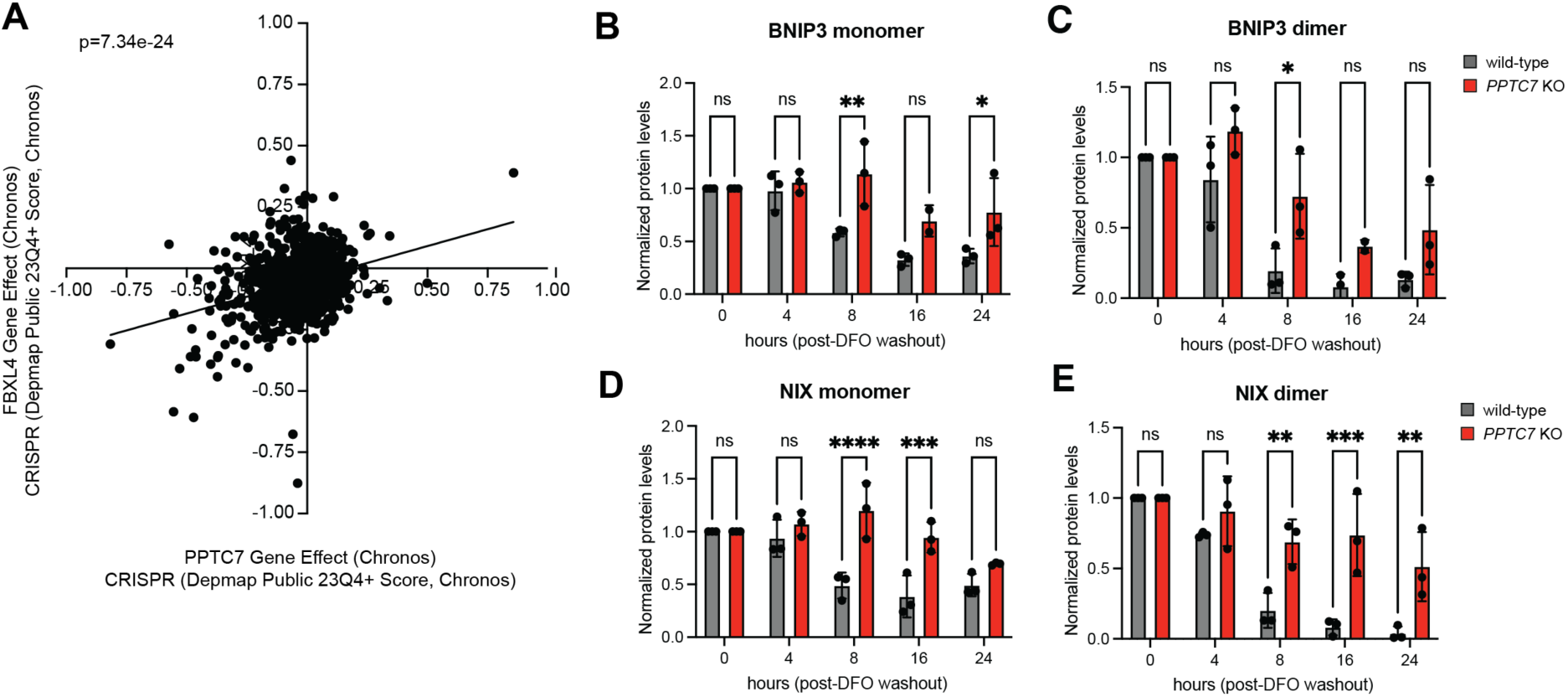
**A**. DepMap essentiality profiles of PPTC7 gene effect (x-axis) and FBXL4 gene effect (y-axis). Linear regression analysis and associated p-value shown. **B**.-**E**. Statistical analysis of BNIP3 and NIX monomer and dimer turnover rate data in wild-type (gray bars) and *PPTC7* KO (red bards) 293T cells shown in Figure 2. ****p<0.0001, ***p<0.001, **p<0.01, *p<0.05, ns = not significant, ordinary one-way ANOVA. Error bars represent standard deviation, data points represent individual biological replicates.

**Supplemental Figure 3:**
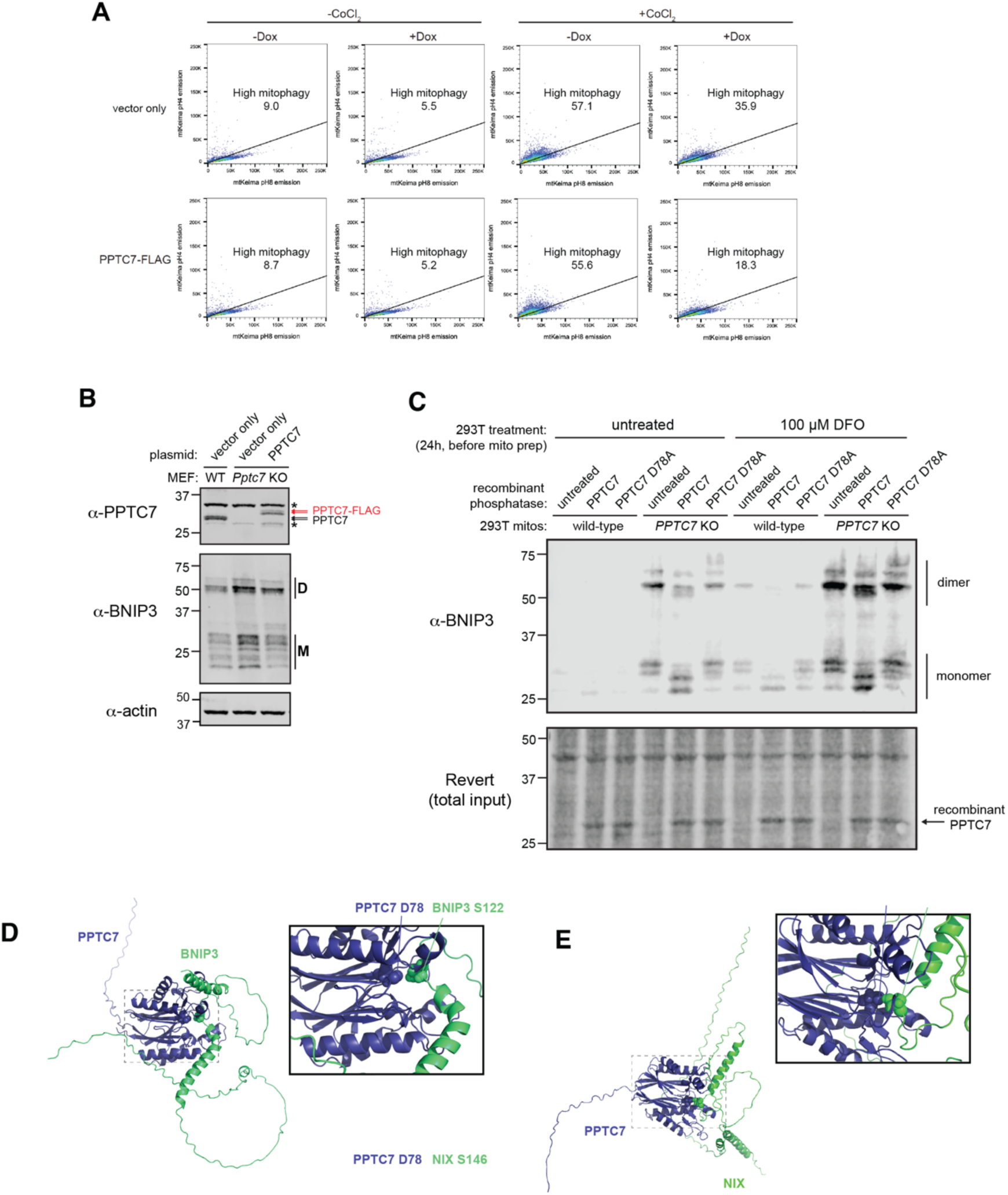
**A**. FACS data shown for data represented in Figure 2D. Cells undergoing high mitophagy are above the diagonal line; percentages indicated in figure. **B**. Western blot of PPTC7 expressed in wild-type MEFs (black arrows), *Pptc7* KO MEFs, or *Pptc7* KO MEFs rescued with human PPTC7 (red arrows). * represents a non-specific band. Basal BNIP3 levels across samples shown below; actin shown as a loading control. **C**. Western blot of BNIP3 in crude mitochondria isolated from wild-type or *PPTC7* KO 293T cells. Mitochondria were left untreated, treated with recombinant PPTC7, or treated with recombinant PPTC7 D78A. Revert stain is shown for loading; equal loading of recombinant proteins can be seen as depicted by arrows. **D**., **E**. AlphaFold2 multimer predictions of the BNIP3-PPTC7 (**D**.) and the NIX-PPTC (**E**.) interaction. Residues of BNIP3 and NIX proximal to PPTC7 D78 highlighted in zoom-in image.

## Methods

### Supplies and reagents

Deferoxamine (DFO) and cobalt chloride were purchased from MilliporeSigma (Burlington, MA). MG132 and polyethylenimine (PEI) was purchased from Fisher Scientific (Hampton, NH). Lipofectamine 3000 was purchased from ThermoFisher (Waltham, MA). Fugene 6 was purchased from Promega (Madison, WI). Multiple plasmids were ordered or generated for this study, including: myc-*Mm*BNIP3-FL in pcDNA3.1, which was a kind gift from Joseph Gordon (Addgene #100796) and V5-miniTurbo-NES in pcDNA3.1, which was a kind gift from Alice Ting (Addgene #107170). pCMV-SPORT6-*Mm*NIX was purchased from Horizon Discoveries. pcDNA3.1-PPTC7-FLAG, pcDNA3.1-FLAG-*Mm*NIX, pcDNA3.1-PPTC7-GFP, pcDNA3.1-PPTC7-OMP25-FLAG, pcDNA3.1-PPTC7-V5-miniTurbo, and pcDNA3.1-ΔMTS-PPTC7-FLAG were generated for this study through standard PCR and restriction-enzyme based cloning techniques. Plasmids for the HeLa FLP-IN TREx system were generated using Gateway cloning. Briefly, constructs were PCR amplified with primers containing attB1 or attB2 sequences. These fragments were incubated with pDONR221 (a kind gift from Julia Pagan) and BP clonase for recombination. Positive constructs were incubated with LR clonase and the pcDNA5/FRT/TO-Venus-Flag-Gateway destination vector, which was a kind gift from Jonathon Pines (Addgene #40999). All cloned plasmids were validated via Sanger sequencing.

### Cell culture and Transfection

293T were acquired from the American Type Culture Collection (Manassas, VA). *PPTC7* KO 293T cells were generated as previously described (Meyer *et al*, 2020) and MEF cells were generated from wild-type and *Pptc7*^-/-^ mice embryos as previously described (Niemi *et al*, 2023). Wild-type, *Pptc7* KO, and *Pptc7/Bnip3/Nix* TKO MEFs were transduced with mt-Keima as previously described (Niemi *et al*, 2023). HeLa FLP-IN TREx cells stably expressing mt-Keima were a kind gift from Dr. Julia Pagan. Cells were cultured in growth media (Delbecco’s Modified Eagle Media supplemented with 10% heat inactivated Fetal Bovine Serum and 1× penicillin/streptomycin). Cells were grown in a temperature-controlled incubator at 37°C and 5% CO_2_. Transient plasmid transfection into 293T wild-type and *PPTC7* KO cells was performed with polyethylenimine (PEI) for 24 to 48 hours. Transient plasmid transfection into U2OS and HeLa cells was performed with Lipofectamine 3000 for 5 hours. Stable plasmid transfection into HeLa FLP-IN cells were performed in the presence of pOG44 in a ratio of 0.5 μg pcDNA5 to 2 μg pOGG44. Transfections were performed with Fugene 6 (Promega) per manufacturer’s directions for 24-48 hours before selection with 400 μg/mL hygromycin B.

### SDS-PAGE and immunoblotting

Cells were lysed with radioimmunoprecipitation buffer (RIPA; 0.5% w/v sodium deoxycholate, 150mM sodium chloride, 1.0% v/v IGEPAL CA-630, 1.0% sodium dodecyl sulfate (SDS), 50mM Tris pH8.0. 1mM EDTA pH8.0 in water) unless otherwise specified. After generating cell lysates, samples were clarified by centrifugation (21,100×*g*) at 4°C for 10 minutes, snap frozen in liquid nitrogen, and stored at –80°C before use. All samples were quantified with the bicinchoninic acid (BCA) assay kit (Thermo Scientific). Lysates were mixed with 5x sample buffer (312 mM Tris-Base, 25% w/v sucrose, 5% w/v SDS, 0.05% w/v bromophenol blue, 5% v/v β-mercaptoethanol, pH6.8) and boiled at 95°C for 10 minutes. Lysates (20-40 μg) were run on SDS-PAGE gels with Precision All-Blue Protein Standards (BioRad) before being transferred onto nitrocellulose membranes. Membranes were incubated with primary antibodies with 2% BSA or 3% nonfat diary milk in TBS-T for periods as indicated above. Primary antibodies used in immunoblotting include: anti-human BNIP3 (Cell Signaling Technology (CST) catalog #44060, dilution 1:1000, 48 hour incubation at 4°C), anti-rodent BNIP3 (CST catalog #3769, dilution 1:1000, 48 hour incubation at 4°C), anti-NIX (CST catalog #12396, dilution 1:1000, 48 hour incubation at 4°C), anti-PPTC7 (Novus catalog #NBP190654, dilution 1:1000, 48 hour incubation at 4°C), anti-HIF1α (CST catalog #36169, dilution 1:1000, overnight incubation at 4°C), anti-β-actin (CST catalog #3700, dilution 1:1000; CST catalog #4970, dilution 1:1000; and Abcam catalog #ab170325, dilution 1:1000; overnight incubation at 4°C), anti-FLAG (Sigma catalog #F1804, dilution 1:2000, overnight incubation at 4°C), anti-V5 (Fisher catalog #PIMA515253, dilution 1:1000, overnight incubation at 4°C), and anti-myc (Fisher catalog #MA121316, dilution 1:1000, overnight incubation at 4°C). Membranes were washed 2-3x with TBS-T for 5 minutes per wash and incubated with corresponding fluorophore-conjugated antibodies for 30 minutes at room temperature. Anti-680 or anti-800 conjugated mouse or rabbit antibodies (LiCOR) were used for detection. Membranes were then washed 2-3x with TBS-T for 5 minutes per wash and were imaged with a LiCOR OdysseyFC instrument using Image Studio software (LiCOR; version 5.2).

### RNA extraction and qRT-PCR

For RNA extraction, the collected cell pellets were processed using Monarch RNA extraction kit (New England BioLabs). The RNA samples were then quantified with nanodrop and normalized before cDNA synthesis (Lambda Biotechnologies). Quantitative PCR analyses with SYBR Green PCR Master Mix (Applied Biosystems) were done on a BioRad CFX-96 Touch Real-Time PCR Detection System controlled by CFX Maestro (ver2.2) on a computer. Primers used were: GAPDH forward 5’-TTCGCTCTCTGCTCCTCCTGTT-3’, GAPDH reverse 5’-GCCCAATACGACCAAATCCGTTGA-3’, BNIP3 forward 5’-GCCCACCTCGCTCGCAGACAC-3’, BNIP3 reverse 5’-CAATCCGATGGCCAGCAAATGAGA-3’, NIX forward 5’-CTACCCATGAACAGCAGCAA-3’, and NIX reverse 5’-ATCTGCCCATCTTCTTGTGG-3’.

### Flow cytometry

For all experiments, wild-type MEFs or HeLa cells (i.e., with no mt-Keima expressed) were used as a negative control and were plated at the same density as other cells in the corresponding experiment. Cells were grown in standard DMEM media (25 mM glucose, 2 mM glutamine, 10% heat-inactivated FBS, 1x penicillin/streptomycin) for 48 hours. Cells were harvested by trypsinization and were resuspended in FluoroBrite media with 0.8% heat-inactivated FBS in 5mL polystyrene round-bottom tubes (Falcon) immediately prior to the flow cytometry experiments. The LSR-Fortessa (BD Biosciences) flow cytometer was used for the experiment, and the flow cytometer was controlled by BDFACSDiva (version 9.0). FSC (488 nm), SSC (excitation 488nm, emission 488 nm), QDot 605 (excitation 405nm, emission 610 nm), and PE-TexasRed (excitation 585 nm, emission 610 nm). The laser intensities for Qdot 605 and PE-TexasRed were changed based on the emission profile of MEF wild-type cells for each experiment and were kept constant throughout the experiment. Cells were gated to select for live cells, single cells, and mt-Keima positive cells sequentially. Once gates were established in a given experiment, they were used for the duration of that experiment. Flow cytometry data were processed with FlowJo (version 10.2.2), and the high mitophagy gate was drawn during analysis.

### Immunofluorescence assay and confocal microscopy

For immunofluorescence assay, cells grown on glass-bottom cover dishes (CellVis) were fixed in 4% paraformaldehyde solution in PBS (15 minutes, room temperature). Fixed cells were permeabilized (0.1% Triton X-100 in PBS), blocked (10% FBS and 0.1% Triton X-100 in PBS), and then incubated overnight at 4°C with the indicated primary antibodies in blocking buffer. Primary antibodies used in immunofluorescence assays include anti-TOMM20 (Abcam catalog #56783, dilution 1:400) and anti-human BNIP3 (CST catalog #44060, dilution 1:100). After three washes (5 minutes each) in PBS, cells were incubated with secondary antibodies in blocking buffer for 30 min. The secondary antibodies used in immunofluorescence assays include donkey anti-mouse Alexa Fluor 647 (Fisher catalog #A-31571, dilution 1:400) and donkey anti-rabbit Alexa Fluor 555 (Fisher catalog #A-31572, dilution 1:400). Cells were subsequently washed three times in PBS prior to imaging. Images were acquired on a Nikon Ti2 microscope equipped with Yokogawa CSU-W1 spinning disk confocal and SoRa modules, a Hamamatsu Orca-Fusion sCMOS camera and a Nikon 60x objective (for Figure 3A-B) or Nikon 100x 1.45 NA (for Figure 4A-B) objective. All images were acquired using a 0.2-μm step size with the spinning disk module, and image adjustments were made with ImageJ/Fiji.

### DFO-induced iron chelation and washout

For immunoblotting experiments to visualize changes of BNIP3 and NIX protein levels upon iron chelation, 293T wild-type and *PPTC7* KO cells were treated with 100 μM DFO for 0-24 hours as indicated in the results section. After the indicated incubation time, the cells were harvested by cell scraping in phosphate-buffered saline (PBS) before cell lysis and immunoblotting analysis.

For qRT-PCR experiments to quantify changes in *BNIP3* and *NIX* transcript level, 293T WT and *PPTC7* KO cells were treated with 100μM DFO for 24 hours, and the cells were harvested by cell scraping in PBS before RNA extracton and qRT-PCR analysis.

For DFO washout experiments, after treating 293T WT and *PPTC7* KO cells with 100μΜ DFO for 24 hours, cells were washed with PBS and was switched to fresh DMEM media (for Figure 2B-F) or with DMEM containing 0.1% EtOH (as vehicle control) or 10 μM MG132 (for Figure 2I) for 0-24 hours as indicated in the results section. After the indicated incubation time, the cells were harvested by cell scraping in phosphate-buffered saline (PBS) before cell lysis and immunoblotting analysis.

For flow cytometry experiments to determine the change of mitophagic flux with DFO dosage, MEF mt-Keima wild-type and *Pptc7* KO cells were treated with 0-100 μM DFO as indicated in the results section for 24 hours before downstream processing for flow cytometry. To understand whether the DFO-induced mitophagy is BNIP3– and NIX-mediated, MEF mt-Keima wild-type, *Pptc7* KO, and *Pptc7/Bnip3/Nix* TKO cells were treated with 100μM DFO for 24 hours before downstream processing for flow cytometry.

### MG132 proteasomal inhibition time course

For MG132 proteasomal inhibition time course experiment done in Figure 2G-H, 293T WT and *PPTC7* KO cells were treated with 0.1% EtOH (as vehicle control) or 10 μM MG132 for 0-6 hours as indicated in the results section. After the indicated incubation time, the cells were harvested by cell scraping in PBS before cell lysis and immunoblotting analysis.

### Analysis of PPTC7 localization relative to BNIP3

U2OS cells were transiently transfected with PPTC7-GFP plasmid. Cells were then passaged into glass-bottom dishes. Cells were allowed to adhere for 12 hours and subsequently treated with freshly prepared DFP (1 mM; Sigma-Aldrich) for 24 hours prior to fixation. Where indicated, cells were then washed with fresh media, and fixed after an additional 4 hours of incubation. Immunofluorescence staining and confocal microscopy were then performed on cells. To determine the co-enrichment of PPTC7 with BNIP3, enlarged foci enriched for BNIP3 signal relative to TOMM20 were manually counted in Fiji by examining single plane images throughout z-series of individual cells blinded to the corresponding PPTC7 image, followed by assessment of whether PPTC7 was co-enriched.

### Analysis of the effect of PPTC7 overexpression on BNIP3 immunofluorescence

For analysis of BNIP3 staining by immunofluorescence, HeLa cells were transiently transfected with cytosolic GFP or PPTC7-GFP plasmids. Cells were allowed to adhere to glass-bottom dishes for 12 hours and treated with freshly prepared CoCl_2_ (500 μM; Sigma-Aldrich) 12 hours prior to fixation. Cells were imaged with identical imaging conditions between experimental replicates. Cells that had GFP signal above an arbitrary threshold that was consistently maintained between experiments were blindly identified and manually categorized as having mitochondrial or diffuse non-mitochondrial BNIP3 signal in Fiji by examining maximum z-projections.

### Analysis of the effect of PPTC7 mutant expression on BNIP3 and NIX total protein levels

HeLa FLP-IN T-REx cells expressing wild-type PPTC7 or various mutants (e.g., D78A) were generated as described above. To induce expression of these constructs, cells were treated with 10 μM doxycycline for 24-32 hours, after which BNIP3 and NIX expression was induced by treating cells with 200 μM CoCl_2_ for 16 hours. Cells were collected, lysed in RIPA buffer, run via SDS-PAGE and western blotted for each protein as described above (see “SDS-PAGE and immunoblotting”).

### Proximity labeling

Wild-type 293T cells were transiently transfected with pcDNA3.1-MCS (empty vector), pcDNA3.1-V5-miniTurbo-NES, or pcDNA3.1-PPTC7-V5-miniTurbo before cells were treated with either water or 100 μM DFO (supplemented in growth media) for another 24 hours. For labeling, DMSO or 250 μM biotin were added to cells, and cells were incubated at 37°C for 30 minutes. Labeling reactions were quenched by washing the cells three times with ice-cold PBS and incubating the cells at 4°C. Cells were then harvested in ice cold PBS before lysed with RIPA, clarified, and quantified via BCA. The lysates were then incubated with 80 μL streptavidin magnetic beads (New England Biolabs) at 4°C overnight. The beads were then washed twice with RIPA buffer, once with Wash Buffer 2 (500 mM sodium chloride, 0.1% w/v deoxycholate, 1% Triton X-100, 1 mM EDTA, 50 mM HEPES, pH 7.5), once with Wash Buffer 3 (250 mM sodium chloride, 0.5% Triton X-100, 0.5% w/v deoxycholate, 1 mM EDTA, 50 mM HEPES, pH 8.1), once with Wash Buffer 4 (150 mM sodium chloride, 50 mM HEPES, pH 7.4), and once with Wash buffer 5 (50 mM ammonium bicarbonate in MS-grade water). The bound species were eluted by adding 20 μL 5x sample buffer, boiling at 95°C for 10 minutes, vortexing for 10 minutes, and boiling at 95°C for 10 minutes. The samples were then run on SDS-PAGE gels, and immunoblotting was performed to probe for BNIP3 and NIX.

### Data/statistical analysis and figure generation

Statistical analysis was performed using Microsoft Excel and/or Prism software (GraphPad, version 10). Supplemental Figure 1A was generated using Seaborn library on Python. A selected list of HIF target genes were curated (Dengler *et al*, 2014) as mined from datasets in mouse *Pptc7* KO heart and liver proteomics (Niemi *et al*, 2019). Figure 2C was generated using AlphaFold2. Supplemental Figures 2C-D were generated using AlphaFold multimer mode; images were generated using PyMOL (Version 2.5.2). Figure 3A was generated using BioRender with an appropriate license. All figures were generated using Adobe Illustrator.

## Notes

### Competing Interest Statement

The authors have declared no competing interest.

